# Reorganization of the Primate Dorsal Horn in Response to a Deafferentation Lesion Affecting Hand Function

**DOI:** 10.1101/818716

**Authors:** Karen M. Fisher, Joseph Garner, Corinna Darian-Smith

## Abstract

The loss of sensory input following a spinal deafferentation injury can be debilitating, and this is especially true in primates when the hand is involved. While significant recovery of function occurs, little is currently understood about the reorganization of the neuronal circuitry, particularly within the dorsal horn. This region receives primary afferent input from the periphery, and cortical input via the somatosensory subcomponent of the corticospinal tract (S1 CST), and is critically important in modulating sensory transmission, both in normal and lesioned states. To determine how dorsal horn circuitry alters to facilitate recovery post-injury, we used an established deafferentation lesion model (DRL/DCL – dorsal root/dorsal column) in male monkeys to remove sensory input from just the opposing digits (D1-D3) of one hand. This results in a deficit in fine dexterity that recovers over several months. Electrophysiological mapping, tract tracing, and immunolabeling techniques were combined to delineate specific changes to dorsal horn input circuitry. Our main findings show that (1) there is complementary sprouting of the primary afferent and S1 CST populations into an overlapping region of the reorganizing dorsal horn, (2) S1 CST and primary afferent inputs connect in different ways within this region to facilitate sensory integration (3) there is a loss of larger S1 CST terminal boutons in the affected dorsal horn, but no change in the size profile of the spared/sprouted primary afferent terminal boutons post-lesion. Understanding such changes helps to inform new and targeted therapies that best promote recovery.

**Significance statement:** Spinal injuries that remove sensation from the hand, can be debilitating, though functional recovery does occur. We examined changes to the neuronal circuitry of the dorsal horn in monkeys following a lesion that deafferented three digits of one hand. Little is understood about dorsal horn circuitry, despite the fact that this region loses most of its normal input after such an injury, and is clearly a major focus of reorganization. We found that both the spared primary afferents and somatosensory corticospinal efferents sprouted in an overlapping region of the dorsal horn after injury, and that larger (presumably faster) corticospinal terminals are lost, suggesting a significantly altered cortical modulation of primary afferents. Understanding this changing circuitry is important for designing targeted therapies.

## Introduction

The dorsal horn of the cervical spinal cord is central to sensorimotor function. All primary afferent input first terminates here en route to higher regions of the central nervous system (CNS). This area also receives direct input from the cortex, by way of the somatosensory corticospinal tract (S1 CST), which is likely to serve a role in the modulation or gating of peripheral sensory information as it enters the cord (Seki et al., 2003; Seki and Fetz, 2012). Following a spinal deafferentation injury to the hand, which we have examined previously (Darian-Smith et al., 2014; Fisher et al., 2018), the affected region of the dorsal horn permanently loses most (>95%) of its normal input from the periphery. This means that the reorganization of affected local (and distant) neuronal circuitry in the early post-lesion weeks and months is critical to the recovery of digit and hand function. Despite its importance in sensory processing, the intrinsic circuitry of the dorsal horn is poorly understood in both the normal or injured state, particularly in primates. We begin to address this here.

We first asked if the greatly diminished primary afferent projection also sprouts following a combined dorsal root/dorsal column lesion (DRL/DCL) in the monkey. Earlier work showed that following a dorsal root lesion alone, the few remaining spared primary afferents sprouted within the cervical dorsal horn on the side of the lesion, presumably to form new functional synapses with postsynaptic targets that had lost their normal input (Darian-Smith, 2004). More recently, we showed that there are some fundamental differences in the CST response between injuries that are purely peripheral (as in the DRL alone), and those involving a central lesion (Darian-Smith et al., 2014; Fisher et al., 2018). Given this work, which showed extensive S1 CST terminal sprouting only when a central dorsal column lesion was involved, we asked if the primary afferent sprouting response was the same or more extensive (or even bilateral) following a central lesion (i.e. a DRL/DCL)?

We next asked what role the S1 CST plays in cervical dorsal horn reorganization following a combined DRL/DCL, at the level of the neuronal circuit. Unlike the M1 CST, S1 CST axons project discretely to the dorsal horn where the terminal territory overlaps with that of the primary afferent input. This relationship supports a role of the S1 CST in gating/integrating primary afferent information, as it enters the cord. The M1 CST is known to have a small ipsilateral projection and to cross the midline (Ralston and Ralston, 1985; Dum and Strick, 1996; Galea and Darian-Smith, 1997; Morecraft et al., 2013), which provides an anatomical framework for terminal bilateral sprouting post-lesion (Rosenzweig et al., 2009; Rosenzweig et al., 2010). However, the source of the S1 CST bilateral terminal labeling is not clear, and is addressed here.

*Finally*, we used an immunofluorescence approach to ask what the primary afferent and S1 CST efferents synapse onto within the spinal dorsal horn. This is the first study to look at changes in this circuitry post-lesion in nonhuman primates. Importantly, we used a species with manual dexterity, and sensorimotor skills similar to humans. In so doing, we model a synaptic relationship between the primary afferent and efferent inputs to the dorsal horn, that is likely to be similar in humans.

We demonstrate for the first time that both peripheral and cortical sensory inputs respond to a small deafferentation lesion by sprouting significantly within the dorsal horn. This occurs over 10 spinal segments, and the two populations show distinct yet complementary responses within a shared neuronal circuitry. Our findings provide a foundation for understanding how new functional connections form after spinal injury to mediate the recovery of hand function.

## Materials and Methods

We collected data from 6 monkeys in this study (4 DRL/DCL and 2 controls). All were healthy young male macaques (*Macaca fascicularis*; 2.96-4.1kg), who were colony bred (Charles River), and housed individually at the Stanford Research Animal Facility. Each had access to four unit cages (64×60×77cm, depth x width x height per each unit) and were housed in a room with other monkeys. The housing area had a 12 hour light/dark cycle and was maintained at a constant temperature (72-74°F).

We adhered to the ARRIVE guidelines, with the exception that only male monkeys were used. This was due to availability, and the lack of any evidence for gender differences with respect to sensorimotor pathways, hand function and CNS recovery following SCI. Throughout the study, animals had freely available water and primate diet, supplemented daily with fresh fruit, vegetables, and a variety of novel foods and drinks. They also had daily enrichment in the form of behavioral training, primate toys, videos and music.

Animal procedures were carried out in accordance with National Institutes of Health guidelines and the Stanford University Institutional Animal Care and Use Committee.

### Experimental sequence and timeline

Table 1 summarizes the experimental details for each animal. The two control animals underwent a craniotomy during which electrophysiological recordings were made in S1 to locate the area receiving input from digits 1-3 (D1-3). Once identified, multiple tracer injections were placed within this region (Figure 1). Five weeks later, the animals were sedated and injections of Cholera Toxin subunit B (CT-B) were made into the finger pads of digits 1-3 on both hands. One week later they were perfused and the brain and spinal cord tissue was collected.

**Table 1.**

| Monkey ID | Lesion side | Postoperative times (wks) |  |  | Tracer injections |  |
| --- | --- | --- | --- | --- | --- | --- |
|  |  | Time: lesion to craniotomy | Tracer transport (wks) |  | S1 | Primary Afferents |
|  |  |  | S1 CST | Afferents |  |  |
| Control |  |  |  |  |  |  |
| 1601 | - | - | 6 | 1 | BDA | CT-B |
| 1702 | - | - | 6 | 1 | BDA | CT-B |
| Lesioned |  |  |  |  |  |  |
| 1402 | Right | 12 | 7 | - | LYD | - |
| 1602 | Left | 12 | 6 | 1 | - | CT-B |
| 1603 | Left | 12 | 6 | 1 | BDA | CT-B |
| 1604 | Right | - | - | - | - | - |

**Figure 1.**
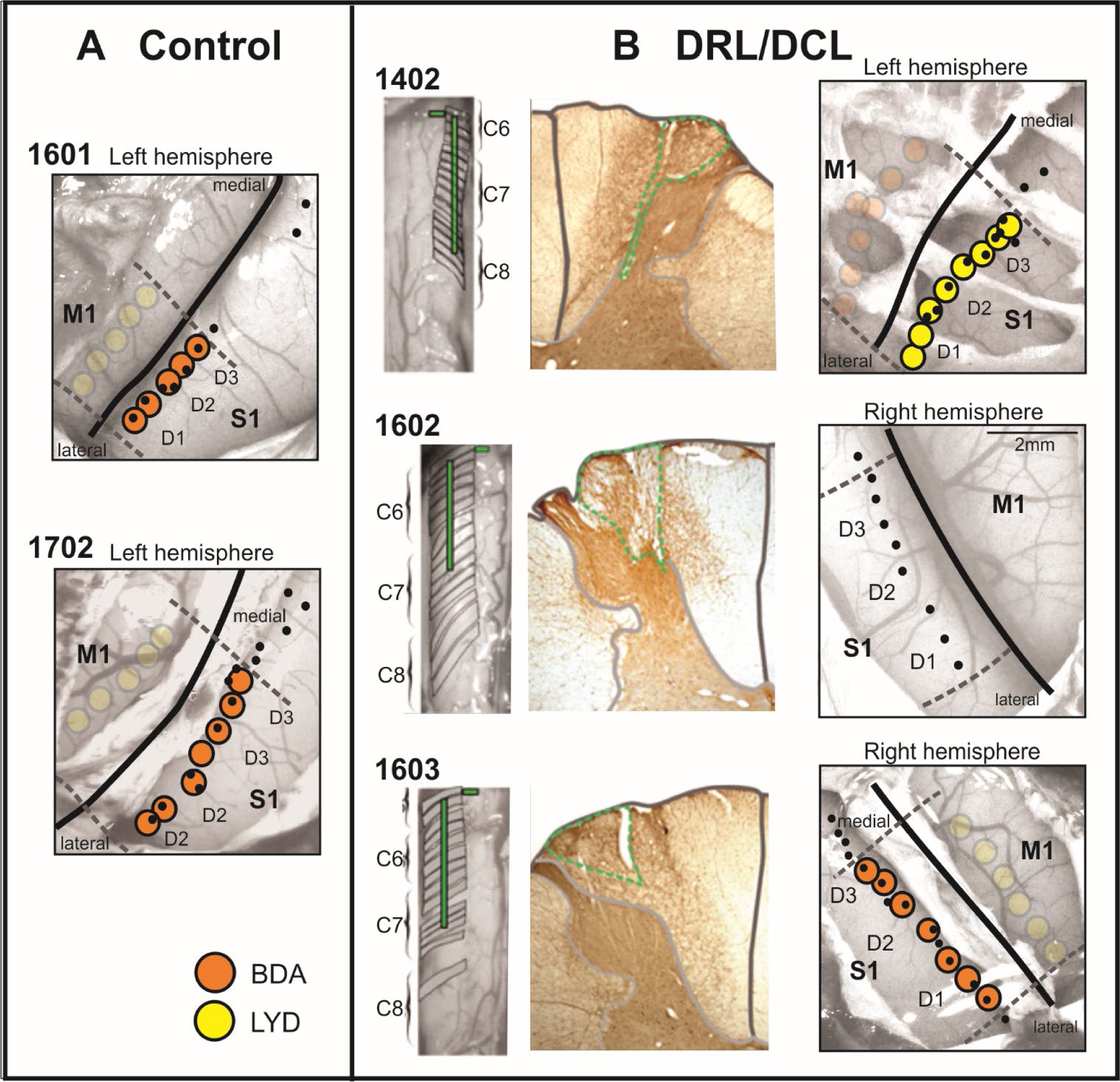
Cortical tracer injection maps and spinal cord lesions (numbers correspond to different animals; see Table 1 for details). ***A***, Schematics showing recording sites (black dots) and injection sites (yellow and orange circles, transparency indicates injections which are not relevant to work discussed in this manuscript) within somatosensory cortex for the control animals. ***B***, Photographs of the exposed spinal cord during surgery showing the lesion locations for the DRL/DCL monkeys. Alongside are photomicrographs of LYD sections close to the DCL core. Green dotted lines delineate the extent of this lesion which was limited to the cuneate fasciculus. Wallerian degeneration (degeneration of fibers distal to site of injury) which expands medially across the dorsal column is a consequence of the DRL. Cortical injection sites are also illustrated for the lesioned animals.

Animals in the lesion group had an initial laminectomy which exposed the spinal cord unilaterally. Recordings were made to identify dorsal rootlets carrying information from digits 1-3; these were then transected and a dorsal column lesion was placed at the rostral border of the D1 territory. Animals recovered for 4-5 months before undergoing a craniotomy and cortical and peripheral tracer injections on the timescale above (Figures 1 & 2).

**Figure 2.**
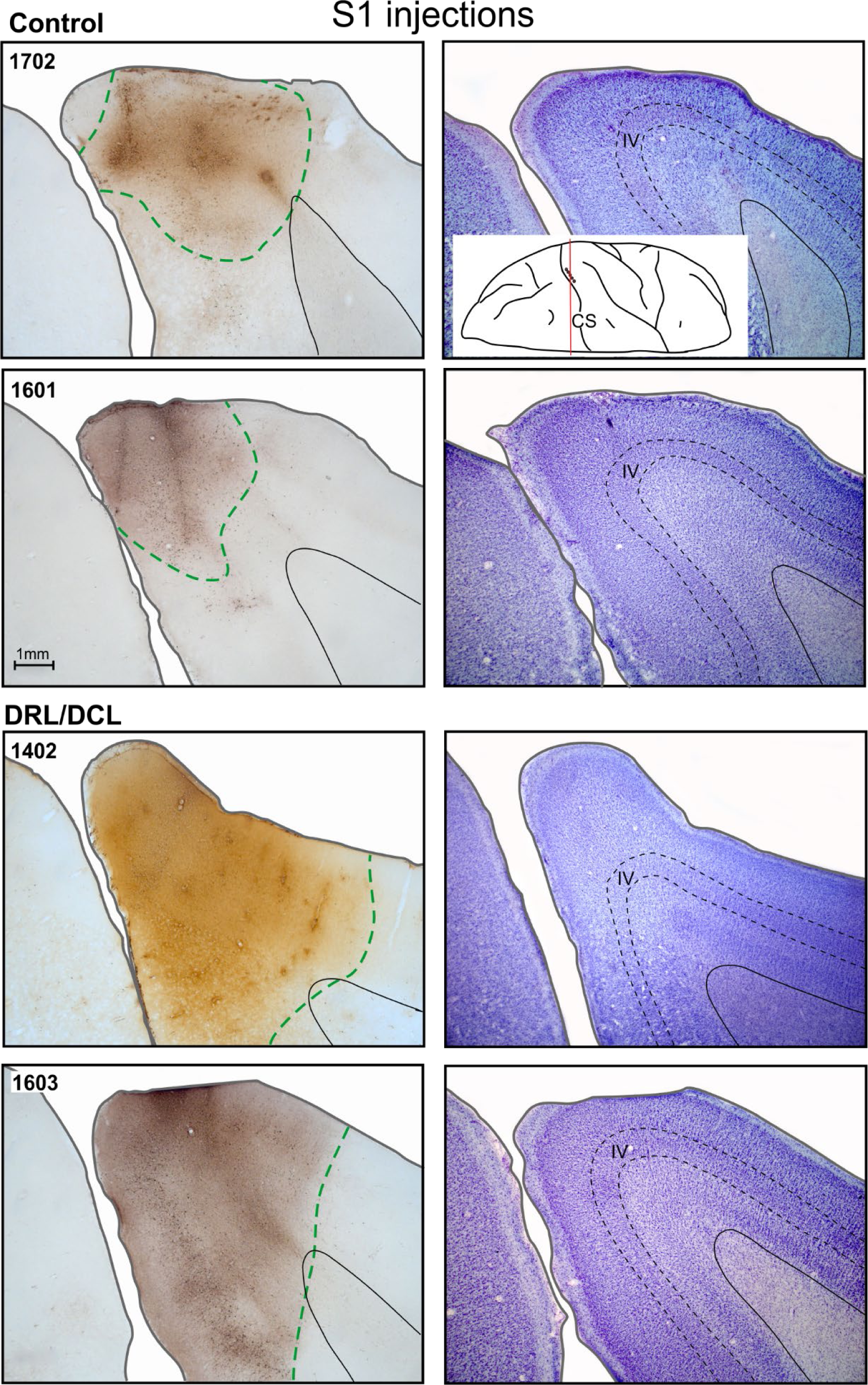
Photomicrographs of tracer injection sites in somatosensory cortex (left column) alongside Nissl stains of neighbouring sections for reference (right column). Green dotted lines designate injection area.

### Behavioral assessment

The monkeys in this study were trained to perform a reach-retrieval task, as has been described in detail elsewhere (Darian-Smith and Ciferri, 2005). Unfortunately, for reasons unrelated to this study, we were unable to obtain complete data sets from these animals. Though we present sample data (see Figures 3 & 4), the behavioral profile described relies heavily on analyses conducted in monkeys not explicitly used in this study. These additional monkeys (i.e. shown in Figure 3 and in a separate report-Crowley et al., unpublished), were the same species, age, and sex as those used in the present study, and all had identical DRL/DCLs affecting D1-D3 in one hand. They also had identical behavioral training routines. As such, the monkeys used here were qualitatively indistinguishable from animals used in other studies in our lab.

**Figure 3.**
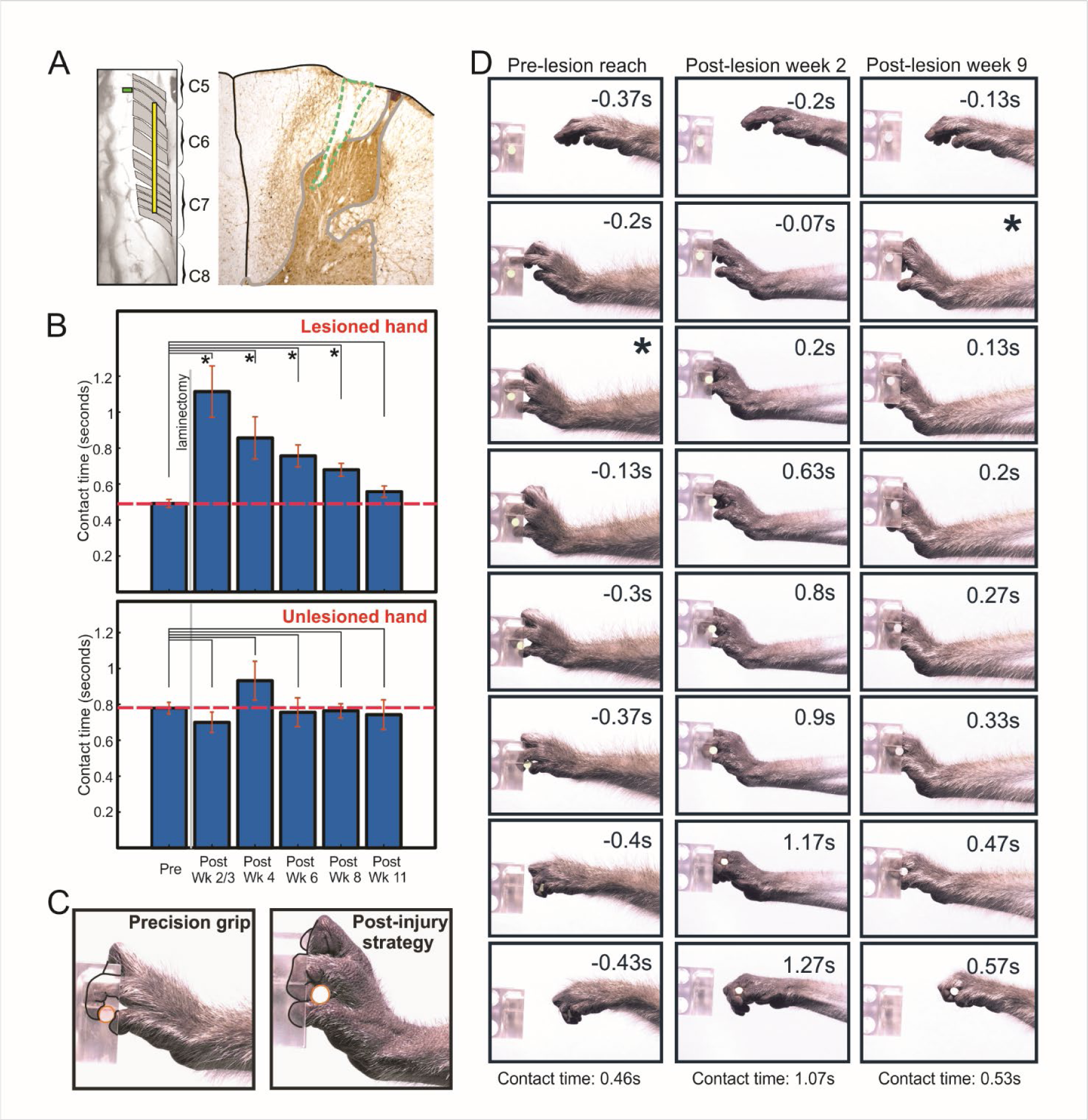
Behavioral data from monkey 1604, for clamp force of 2.0 Newtons. **A**, shows images of the laminectomy with the DRL (left) and DCL placements for this animal. **B**, shows graphs of the mean contact times obtained for the lesioned and unlesioned hands. Bars represent S.E.M.s, and asterisks indicate significant datapoints (P<0.0005). **C**, Stills showing the dominant retrieval strategy used pre- and post-lesion. **D**, shows frame sequences for typical pre- and post-lesion trials. Prior to the DRL/DCL, the monkey retrieved the pellet using a precision grip strategy with the tips of digits 1 and 2 (column 1). In the early weeks post injury (column 2), the sensory deficit resulted in inaccurate digit placement and force application, which caused a significant increase in contact time. By 9 weeks, the monkey had developed a successful compensatory strategy which restored speed, but not the precision grip (column 3). In each series, the star symbol indicates the point at which contact is made with the pellet.

**Figure 4.**
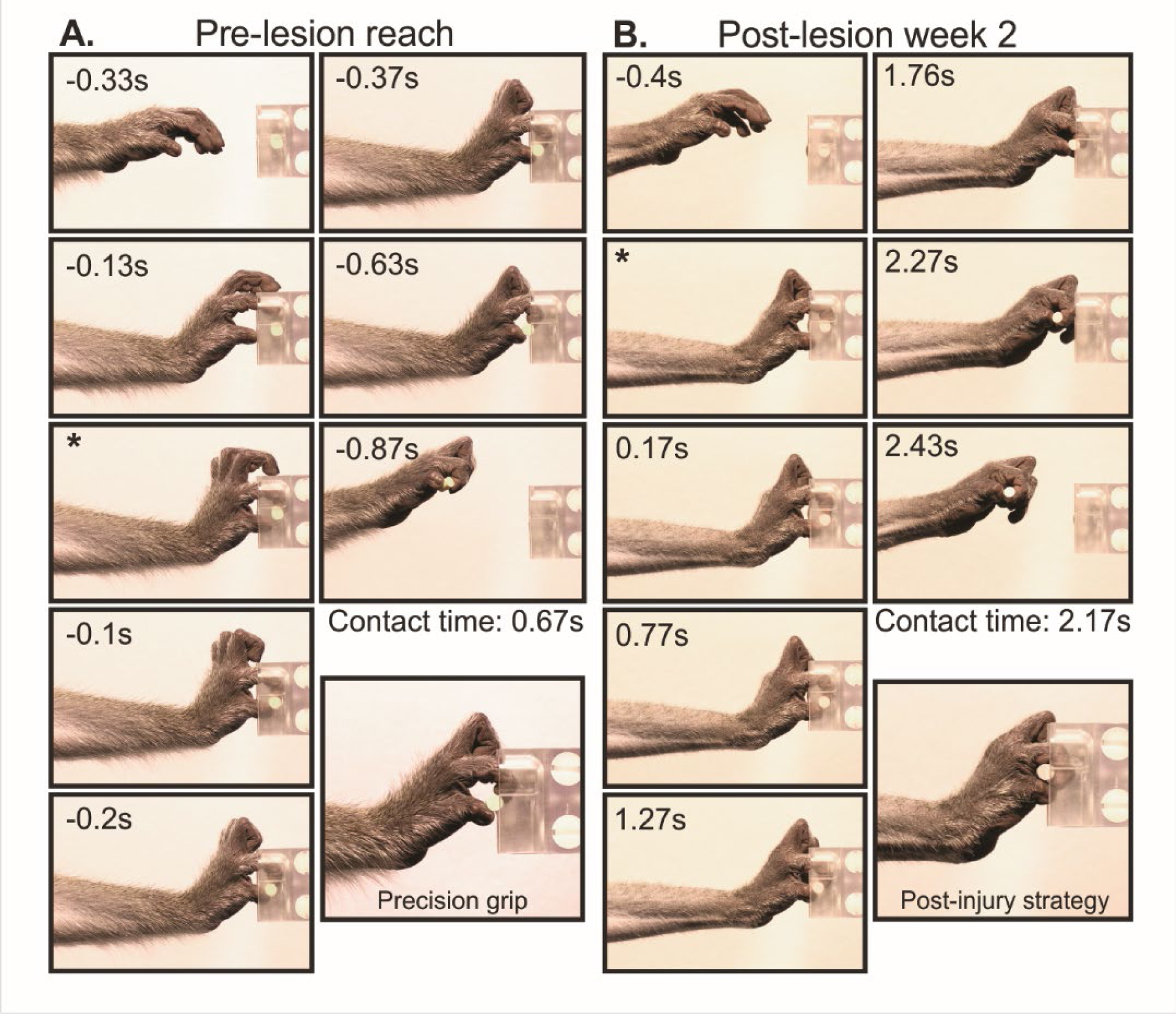
Behavioral data from monkey 1603. Each sequence shows frames from a reach-retrieval trial (2 Newton force) and a typical grip strategy either before or after DRL/DCL (1.5 Newton force). Note the increased contact time post-lesion and the inaccurate placing of the pellet relative to digits 1 and 2. Star symbols indicate time when contact is made with the pellet.

#### Reach-retrieval task

The task involved a natural movement, therefore monkeys learned quickly and performed at a highly consistent speed within 4-6 weeks. Briefly, monkeys were trained to sit in a plexiglass box and reach through a window (located on either the right or left depending on the hand being tested), to retrieve a candy pellet held in one of 4 clamps. Each clamp held the pellet at one of 4 different forces (0.5, 1, 1.5 or 2 Newtons), and clamps (which were visually indistinguishable to the monkey), were presented pseudo-randomly to the monkey. Training sessions were filmed using a high shutter speed (30 fps at 1/500s) digital camcorder (Canon XF200) and data was analysed offline using Edius Pro 9 (Grass Valley) software. We measured the time interval between first contact with the target pellet and its successful displacement from the clamps, and refer to this measurement as ‘contact time’ throughout the manuscript.

### Surgery

For all surgical procedures, animals were sedated with ketamine hydrochloride (10mg/kg), and then maintained under gaseous anesthesia (isofluorane, 1-2% in/1% O_2_), using a standard open circuit anesthetic machine. Atropine sulphate (0.05mg/kg), buprenorphine (0.015mg/kg) and the antibiotic cefazolin (20mg/kg) were administered initially, in addition to a bupivicaine line block of the surgical site prior to the first incision. Normasol-R was infused (i.v.) throughout surgery to maintain fluid balance. For craniotomy procedures, dexamethasone (0.25mg/kg) was also given at the start to minimise brain oedema.

Monkeys were kept warm using a thermostatically controlled heating pad (circulating water) and a Bair Hugger warm air blanket. Physiological signs were carefully monitored throughout surgery to ensure a deep, stable anesthesia (i.e. blood pressure, heart rate, pulse oximetry, capnography, and core temperature).

Buprenorphine (0.015mg/kg) was administered following surgeries in all animals as a post-operative analgesic and monkeys were returned to their cages for recovery. Within an hour, animals were awake and alert. Oral meloxicam (0.1mg/kg) was administered for 3-5 days post-surgery, as needed. Buprenorphine (0.05mg/kg, oral route) was also given as indicated for additional analgesia in the days following surgery.

#### Laminectomy & DRL/DCL lesions

Animals in the DRL/DCL group first underwent a laminectomy to expose spinal segments C5-C8. Lesions were then guided by electrophysiology since the segmental organization of afferent inputs is highly variable both within and between animals, and cannot be predicted by the distribution of the myotome (Dykes and Terzis, 1981).

##### Dorsal root recordings

The dura was resected on one side and a tungsten microelectrode (1.2-1.4mΩ at 1 kHz; FHC) was lowered vertically into dorsal rootlets. Extracellular recordings were made from axons within each fascicle to produce a microdermatome map (Darian-Smith et al., 2000). Cutaneous receptive fields were mapped using hand manipulation, a camel hair brush and Von Frey hairs. Receptive fields were classified as cutaneous if a stimulation force ≤ 2.0g evoked a response. For higher stimulation forces or where responses could only be evoked with joint movement, the receptive field was classified as deep. If there was uncertainty about the nature of a receptive field, this was noted and for the purpose of making the lesion, it was considered cutaneous.

##### DRL/DCL procedure

For the DRL component of the injury, only rootlets with detectable cutaneous receptive fields on the thumb, index and middle fingers, were cut (using iridectomy scissors). Rootlets were cut at two sites to leave a gap across which the nerve could not regenerate. This procedure was as described in previous studies (Darian-Smith and Brown, 2000; Darian-Smith et al., 2013; Darian-Smith et al., 2014).

The DCL was made at the rostral border of thumb input to the spinal cord in order to ensure that the two lesions (DRL and DCL) were functionally comparable across monkeys. We used a micro scalpel blade (Micro-Scalpel, Feather, 15^0^) which was marked 2mm from the tip to guide the depth of the lesion, and only the cuneate fasciculus was targeted.

Once the lesions were completed, the dura was replaced without suturing, and the overlying tissues sutured in layers and skin closed. Tissue glue (3M Vet bond) was applied externally for added security.

#### Craniotomy

All animals received a craniotomy (~1 cm^2^) over the ‘hand’ region of the sensorimotor cortex (in the hemisphere contralateral to the lesion for DRL/DCL animals).

##### Cortical recordings

The somatosensory cortex (areas 3b/1) was mapped using extracellular recording, as described elsewhere (Darian-Smith and Brown, 2000; Darian-Smith et al., 2013; Darian-Smith et al., 2014). This allowed us to determine where input from the partially deafferented digits was localized. Tracer injections were then made into the D1-D3 region of S1 (see Figure 1). Once tracer injections were complete, the bone flap was secured in place using bone wax and Vetbond adhesive, and the overlying incision closed.

### Anterograde Tracers

#### Cortical injections

Separate injection series were made of either Biotinylated Dextran Amine (BDA, 15% aqueous (sterile), Sigma B9139) or Lucifer Yellow Dextran (LYD, 15% aqueous (sterile), ThermoFisher D1825) into S1 (D1-D3 region; see Figure 1 for details of injections in each animal). These were made using a constant-pressure Hamilton syringe mounted in a micromanipulator, with a glass micropipette (diameter ≤ 30µm) attached using Araldite Rapid. Injections (0.3µl) were made at a depth of 0.8-1mm, and the syringe kept in place for 2 minutes. Six to seven weeks were allowed for transport.

#### Digit pad injections

One week prior to perfusion, animals were sedated with ketamine (10mg/kg) and Cholera Toxin subunit B (CT-B, 10 µl, Sigma C9903, 1% aqueous (sterile)) was injected subcutaneously into the distal and middle finger pads of digits 1-3 of both hands, using a Hamilton syringe.

### Perfusion and tissue processing

Animals were sedated with ketamine, intubated, and transferred to isoflurane anesthesia. A lethal dose of sodium pentobarbital (Beuthanasia, i.v.; minimum 100mg/kg) was given and they were perfused transcardially with heparinised saline followed by 4% paraformaldehyde (in 0.1M phosphate buffer solution). The brain and spinal cord were removed and postfixed for 24 hours, before transfer to 20% sucrose (in 0.1M PB) for cryoprotection. They were then dissected, blocked, flash frozen using isopentane and stored at −80C. Tissue was sectioned using a freezing microtome (brain – sagittal plane, 50µm thickness; spinal cord – transverse plane, 40µm).

To visualise BDA, free floating sections were blocked for endogenous peroxidases (Bloxall, Vector labs, SP-6000), washed in 0.1M PB with 0.1% triton, incubated in ABC (Vector, PK-6100, 1hr, room temperature), washed again, and reacted with nickel intensified 3,3’-diaminobenzidine (DAB) with urea peroxidase (Sigmafast, D0426, Sigma Aldrich) for 5-12 minutes.

LYD was visualized immunocytochemically. Sections were blocked (Bloxall, Vector labs), washed with 0.1% triton-X 100 in Tris buffered saline (0.1M TBS-TX), and incubated with anti-Lucifer yellow made in rabbit (Invitrogen; A5750; 1:200 dilution in 2% normal goat serum; 48-60h at 4°C). Tissue was then washed (TBS-TX) and incubated with biotinylated anti-rabbit (Vector, BA-1000; 1:200; 24h). After rinsing, avidin biotin was added to attach peroxidase (ABC kit; Vector, PK-4000) and sections reacted with DAB and urea peroxidase (Sigmafast, D4293, Sigma Aldrich) until the reaction product was clearly visible.

For CT-B processing, sections were first blocked for endogenous peroxidases (Bloxall), washed (0.1M TBS-TX), and then treated with 10% horse serum (2h, at room temperature). Sections were then incubated in anti-CT-B made in goat (List Biological #703; 1:4000 dilution in TBS-TX) and 3% horse serum for 2-3 days at 4°C. This was washed out and replaced with ImmPRESS reagent (HRP anti-goat IgG; Vector labs, MP-7405; 2h, room temperature). Following a series of washes, VIP substrate (Vector labs, SK-4600) was the chromagen used to visualise CT-B terminals. We took care not to dehydrate these slides since VIP is soluble in alcohol; they were instead transferred directly to xylene after drying and immediately coverslipped.

For immunofluorescence, protocols followed a set pattern regardless of the antibodies used. Sections were washed initially in PBS with 0.5% triton-X (PBS-TX) and blocked with a 10% solution of normal serum in PBS-TX (2h, room temperature). Primary antibodies (anti-NeuN, MAB377, Millipore, 1:100; anti-calbindin, CB38, Swant, 1:5000; anti-CT-B, #703, List Biological, 1:4000; anti-VGLUT1, AB5905, Millipore, 1:5000) were diluted in PBS-TX with 3% normal serum and sections were incubated in this cocktail for 48-60 hours at 4 ºC. Following multiple washes, sections were exposed to secondary antibodies in PBS-TX for 30 minutes at room temperature (conjugated secondaries either from Invitrogen or Jackson ImmunoResearch). If BDA was required, sections were subsequently washed and incubated in either ExtrAvidin FITC (E2761, Sigma Aldrich, 1:200) or CY3 (E4142, Sigma Aldrich, 1:200) in PBS-TX overnight. If perineuronal nets were being visualised, sections were exposed to Wisteria Floribunda Lectin (WFA; Vector labs, FL-1351; 1:500 dilution) for 24 hours at the end of the protocol. Final washes were with PBS. Sections were then mounted using 0.5% gelatin, and air dried for 30-45 minutes before coverslipping using antifade mounting media (Prolong Diamond Antifade Mountant, Invitrogen).

### Light microscopy, terminal field tracing, and bouton size analysis

Terminal distributions of BDA, LYD and CT-B labeled axons in the spinal cord were mapped using Neurolucida software in combination with a Lucivid projection (all MBF Bioscience). This was done for sections spanning C1-T2 at a frequency of 400-1600µm (depending on the projection under examination). Contours were drawn around the mapped boutons to outline the distribution territory in each section and the area of this was calculated. Outlying boutons were not included if they were few in number (<5) since they constituted less than 1% of the total population (Darian-Smith et al., 2014; Fisher et al., 2018).

We also carried out an analysis of bouton size. To do this, we selected multiple sections from C5-6, around the lesion site in DRL/DCL animals and equivalent area in controls. We took high magnification (x100 oil objective) images (throughout the Z plane) of the medial dorsal horn (region of peak S1 CST & primary afferent label) using a Zeiss Axiocam 503 color camera and viewed them using Zen Pro software (Zeiss). We set criteria to measure only boutons which were clearly in focus and not obscured by neighbouring boutons/axons. Then we used an ROI function within the software to determine the area of each bouton.

### Confocal microscopy

After processing for 3-4 antibodies, sections were viewed using a Nikon A1R confocal microscope. The region of overlap between primary afferent and S1 CST inputs was located and this region scanned to create a z stack at high magnification (x40/x60 oil objectives). Images were saved for analysis offline, and colabeling determined using orthogonal viewing, and 3D rotational software (Nikon Elements). This allowed us to zoom, crop, and rotate a region of interest rapidly for any-angle visualization.

### Experimental design and statistical tests

Where possible, we used a repeated measures approach to control for systematic variation as described in previous work (Darian-Smith et al., 2014; Fisher et al., 2018). This greatly increases the statistical power in studies which are subject to small numbers of animals. All such analyses were performed in JMP Pro 14 and SAS 9.4.

#### Behavioral analysis

Data was analysed for monkey 1604 (Figure 3) who was trained at least 3 times per week during the testing period. Baseline performance was calculated by pooling data across 4 weeks of pre-lesion training. Following the DRL/DCL, data from each week was treated independently to highlight the initial deficit and gradual recovery. Each post-lesion week was tested for significance against baseline data using two sample t-tests. We did not use a repeated measures approach for this analysis since it involved a very limited dataset.

#### Bouton analysis

Since there appeared to be a visual difference in the size of primary afferent and S1 CST terminals (see Figure 5C), we sought to test whether their mean area and the shape of the size distributions differed statistically. Doing so presented several challenges. *First*, non-normal distributions can appear to change their shape with the mean, when in fact the underlying shape parameter may be unchanged. This is true of Poisson and lognormal distributions, both of which are more likely candidates than the normal distribution. *Second*, while methods exist for comparing the shape of two distributions, we wanted to test for interaction effects (i.e. that differences in the shape of the bouton population distribution between S1 CST and afferent neurons were themselves different in control and lesioned animals). We therefore adopted a novel approach to this problem.

**Figure 5.**
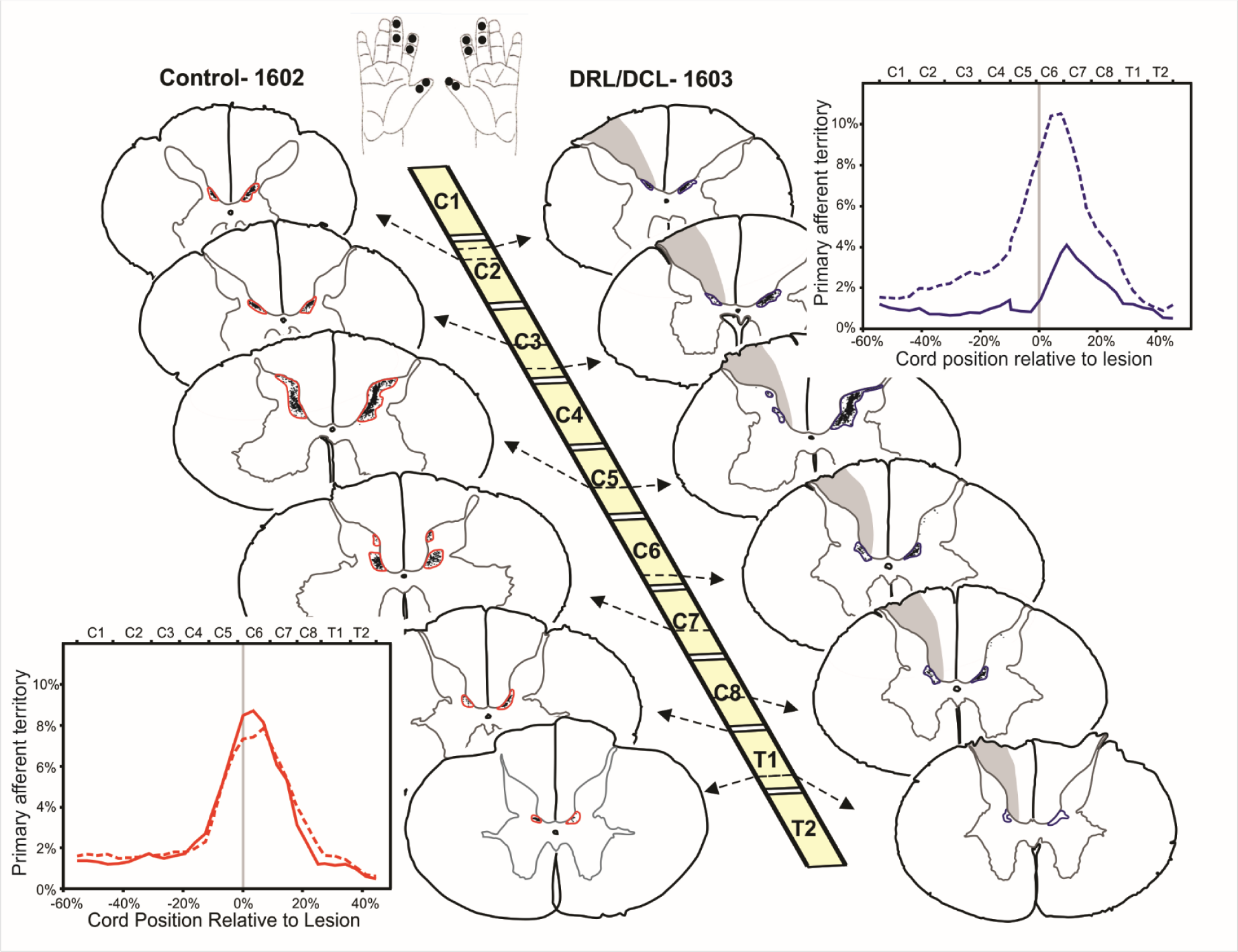
Representative sections through the spinal cord illustrating the primary afferent terminal distribution territory for a control (left) and DRL/DCL monkey (right). The top schematic indicates the location of CT-B injections in the glabrous skin of the digits (black dots). Grey shading indicates regions of Wallerian degeneration following the lesion. Accompanying graphs show terminal territories as a % of grey matter area, for each side of the cord, from C1-T2. Lines are smoothed using a quadratic weighted travelling mean. Grey lines indicate either the location of the DCL in the lesioned animal or the relative cord position in the control. There was no significant difference in the distribution profiles on the two sides of the cord in control animals. By contrast, in monkey 1603, and all lesioned monkeys, the terminal territory was significantly smaller ipsilateral to the lesion.

We first tested the distribution of the bouton area, subdivided by treatment (control or lesion) and bouton type (S1 CST or afferent), against candidate distributions. The normal distribution was consistently the worst fit, and lognormal the best. We therefore log-transformed the bouton areas, and assigned each bouton to a size class defined in log-units. As expected, the resulting distributions were now normally distributed. The resulting counts were approximately poisson distributed, and therefore were square-rooted to achieve a normal distribution for analysis. We then proceeded with a REML mixed-model repeated measures analysis, where size class was treated as a quadratic regression, subject was nested within treatment, and crossed with bouton type. Suitable interactions were included to test whether the shape of the quadratic curve differed for each subpopulation of boutons. None of these interactions were significant, and were pruned from the model following best practice for general linear models and polynomial regression (Grafen and Hails, 2002). The resulting quadratic equations were solved and the peak of each curve were tested against each other using Bonferroni-corrected planned contrasts.

## Results

Five monkeys were used in the central analysis of this study, and the details of their lesions, cortical maps and injection placements are provided in Figure 1. The size and location of the dorsal root and dorsal column lesions was similar across animals, and their DCLs were localized to the cuneate fasciculus component of the dorsal column (Figure 1b). The somatosensory cortex (Areas 3b/1) was mapped to locate the region of digit representation and tracer injections were made only into the D1-D3 region (Figure 1 dotted grey lines). Injections were reconstructed (Figure 2) and did not contaminate neighboring cortical regions or the underlying white matter. Injection volumes were not calculated but were comparable across animals (in terms of volume, injection number and placement), and were similar to those already published (Darian-Smith et al., 2014; Fisher et al., 2018).

### Behavioral observations following a DRL/DCL

No complete data sets were obtained from the monkeys used in this study. However, given the qualitative findings for the monkeys in this study, as well as complete data sets from an additional 4 animals with the same DRL/DCL (Crowley et al., unpublished; see also data from monkeys with a DRL only, Darian-Smith and Ciferri, 2005), we are confident that the behavioral profiles were consistent across animals. All monkeys showed an initial deficit in a precision grip task requiring tactile feedback, and a subsequent recovery of function during the early post-lesion weeks. Representative examples of the behavioral consequences of DRL/DCL are shown in Figures 3 and **4**. Monkeys initially performed the retrieval task using a precision grip involving the opposition of the tips of digits 1 and 2 (Figure 3 C, D; Figure 4). Following the DRL/DCL, retrieval was clumsy, and while they were able to pre-shape the hand normally, they were unable to accurately locate the target using a digit tip precision grip (Figure 3C, Figure 4), or to apply the appropriate force to facilitate retrieval. This meant that contact times were initially significantly longer (Figure 3B, D; Figure 4). Over a recovery period of 5-8 weeks, they developed effective grip strategies, which allowed them to perform the task at pre-lesion speed. This was a functional recovery/compensation, but not a complete restoration of the original strategy.

More detailed behavioral analyses in a different cohort of monkeys (all with equivalent lesions), are the subject of a separate report (Crowley et al., unpublished). All DRL/DCL animals (targeting D1-D3), without exception, have followed a similar deficit and post-injury recovery of hand function.

### Spared primary afferents from the deafferented digits sprout following a DRL/DCL

In control animals, primary afferents carrying sensory information from digits 1-3 terminated within the dorsal horn from spinal segments C1 through T2, with the greatest input observed in segments C5-8. This rostrocaudal spread across 10 segments was far greater than has been previously determined or reported, and is illustrated in Figure 5. For all lesioned monkeys, the primary afferent distribution territory was significantly reduced in the dorsal horn on the side ipsilateral to the lesion, compared with the opposite side, and control animals. An earlier study that looked at primary afferent input following a DRL alone (Darian-Smith, 2004), showed almost no input (conservatively estimated at <5% of the original input), to the ipsilateral cord at 1-2 weeks following the lesion. Given that the current lesion also involved a DRL, combined with an additional central DCL, the assumption can be made that there was a similar near complete removal of input after the DRL/DCL in this study. This means that the ipsilateral distributions observed, despite being considerably reduced, actually reflect terminal sprouting of the spared afferent population on the side of the lesion 4-5 months post-injury. Since earlier work (Darian-Smith et al., 2014; Fisher et al., 2018), showed that the inclusion of a central injury induces bilateral CST sprouting in the cord, we asked in this study if the contralateral primary afferents might also be induced to sprout. However, we found no evidence of any afferent terminal sprouting in the contralateral cord, so the factors leading to bilateral growth of S1 CSTs did not affect the primary afferents in the same way.

The most consistent part of the afferent projection from C1-T2 was observed in the deeper layers of the dorsal horn (Rexed laminae IV-VI). This region receives S1 CST input, contains pre-motor interneurons and is an area that gives rise to the spinothalamic tract. Hantman and Jessell (2010) described a gating circuit for mouse hindlimb proprioceptors in a similar region of the thoracic cord (Clarke’s column). This input extent has not been described previously in the monkey, possibly because most earlier studies focused on inputs to the superficial layers, and cut the cord in the horizontal plane (Florence et al., 1988, 1994; Qi et al., 2016), making it difficult to identify labeling in deeper Rexed laminae.

### S1 CST fibers never cross the midline

Projection patterns for S1 CST in both the control and lesioned state are shown in Figure 6, and were mapped in section series from 4 monkeys (controls =2, lesioned =2; mean no. of sections mapped per monkey = 57). Consistent with previous work (Darian-Smith et al., 2013; Darian-Smith et al., 2014), the S1 CST terminal field expanded following the DRL/DCL, from the normally restricted dorsal horn input zone (see control animal in Figure 6), to a greater proportion of the dorsal horn as well as part of the intermediate region. This expansion occurred throughout the spinal cord, from C1-T2. However, it is important to note that unlike the M1 CST projection, which is known to have a small (~2%; Morecraft et al., 2013) ipsilateral and crossover component, S1 CST fibers were never observed to cross the midline to terminate on the side contralateral to the lesion.

**Figure 6.**
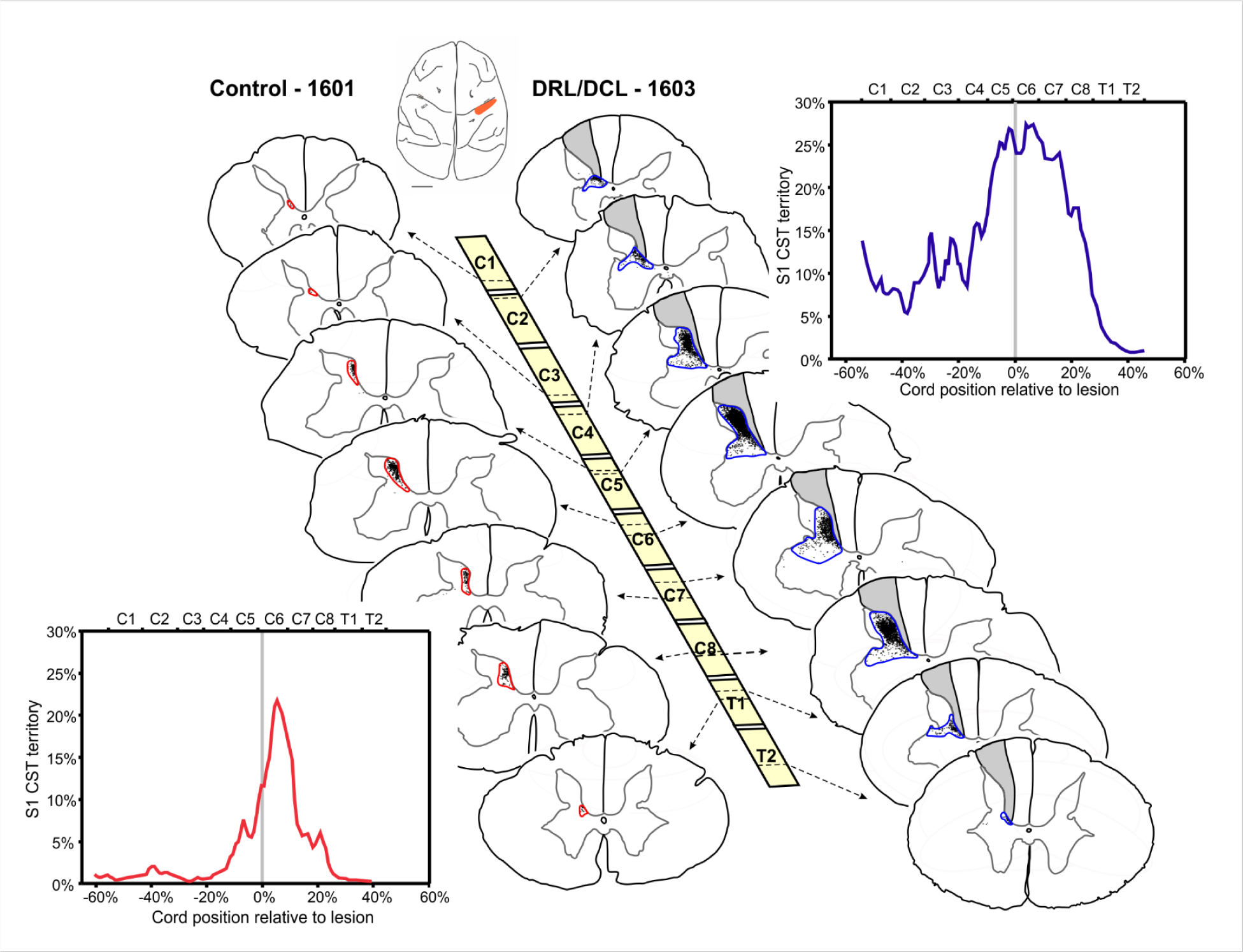
S1 corticospinal terminal labeling in segments C1-T2 of a control animal (left) and following DRL/DCL (right). Terminal areas are outlined in either red (control) or blue (DRL/DCL) and the lesion is always shown on the left side of the cord (grey shading). Note that there is no label contralateral to injections so S1 CST fibers do not cross the midline either in the healthy state or after injury. Inset graphs show distribution profiles for the individuals shown. The lines are smoothed using a quadratic weighted travelling mean. Grey lines indicate the location of the DCL in the lesioned animal or the relative cord position in the control.

### Overlapping populations of afferents and efferent terminals in the dorsal horn support a shared circuitry

Figure 7 shows the relative location of both S1 CST and primary afferent terminals within the dorsal horn at C5/6. On the uninjured side, the normal distribution of primary afferent terminals was visible in the superficial layers (Figure 7A, B, D), whereas they were sparsely present on the injured side despite the sprouting of spared fibers (yellow arrows, Figure 7C). Where BDA is visible (on the injured side), the afferent and CST populations overlay almost completely (see also Figure 8B), so their proximity supports an overlapping functional role.

**Figure 7.**
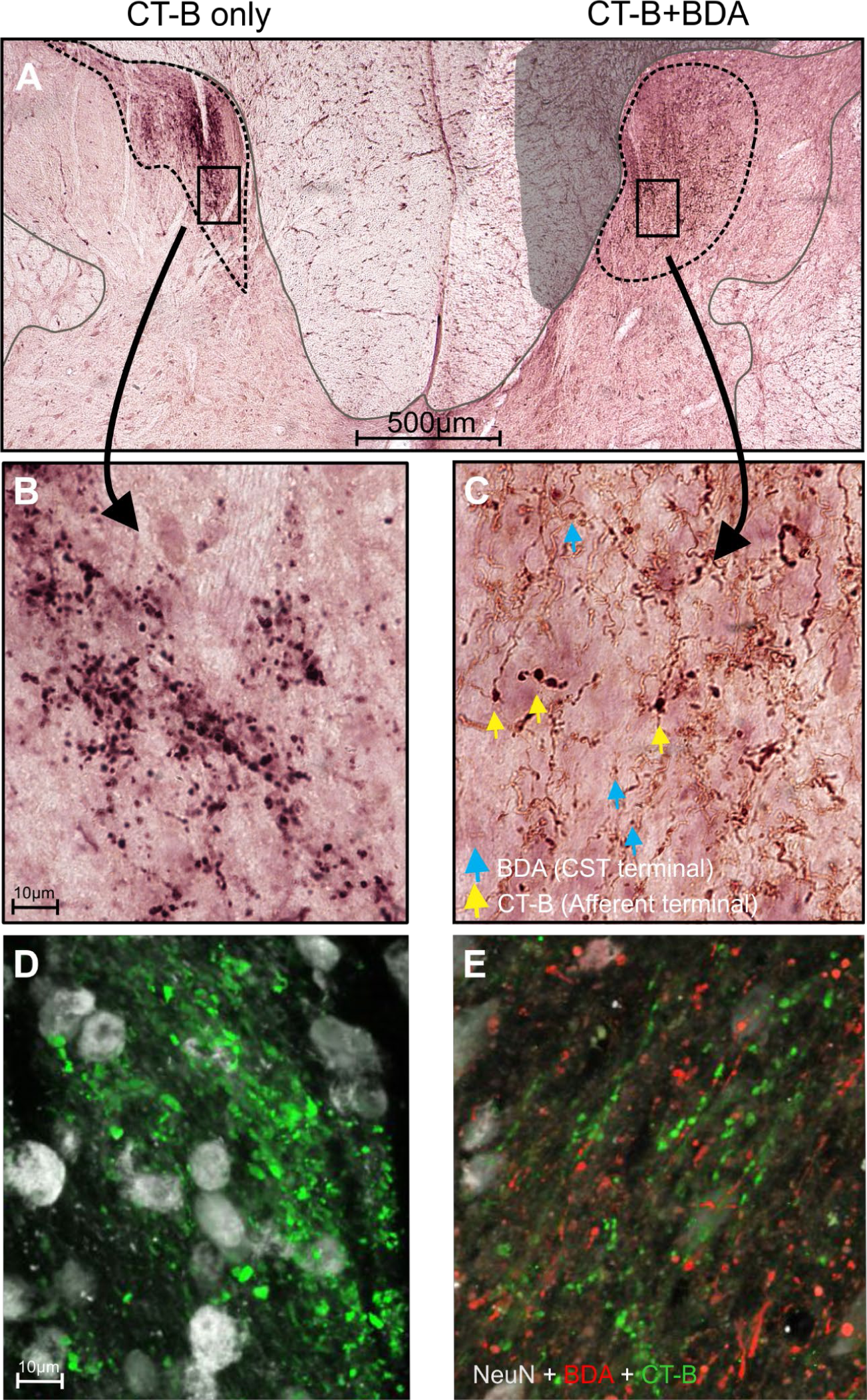
Photomicrographs from a section on the border of C5/6 stained for BDA (S1 CST) and CT-B (primary afferents) in a DRL/DCL monkey (1603). ***A***, Low magnification image showing afferent labeling only (VIP - purple/black) on the left side, and combined afferent and S1 CST labeling (DAB - brown) on the right (lesioned) side. Lesion induced degeneration in the dorsal column is indicated by the gray shaded area. ***B, C***, shows higher magnification images from ***A***. Where BDA and CT-B overlap, S1 CST terminals (blue arrows) and the larger, darker primary afferent boutons (yellow arrows), are indicated. ***D, E***, Adjacent sections processed for immunofluorescence showing distribution of BDA (red) and CT-B (green) terminals. Cells labelled with NeuN are white. Note that many neurons in the region of terminal labeling are NeuN-.

**Figure 8.**
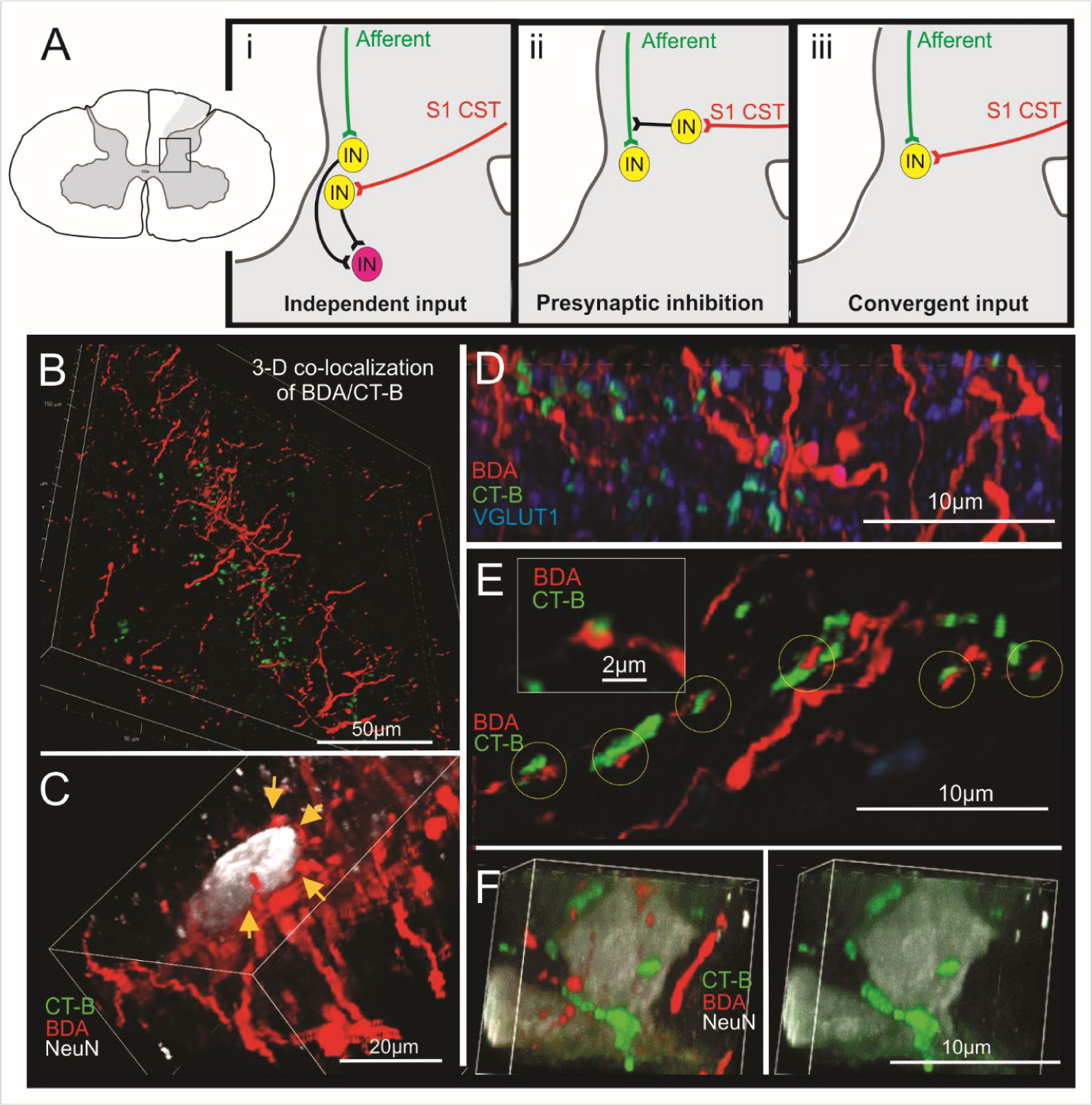
Schematics showing possible mechanisms for sensory integration within the dorsal horn (***A***), as well as supporting confocal images of primary afferent (CT-B) and efferent (BDA=S1 CST), co-localization (***B-F***). ***Ai***, This example shows an indirect route for sensory integration whereby afferent and S1 CST inputs initially contact separate interneuron populations. In ***Aii***, there is gating of primary afferents by S1 CST fibers, and in ***Aiii***, convergence of afferent and S1 CST inputs on to the same postsynaptic target provides a route for direct integration of sensory information within the spinal cord. ***B***, shows the 3-dimensional spatial overlay of the primary afferent (green) and S1 CST (red) terminal populations within the reorganized dorsal horn. ***C***, supports Ai, where only S1 CST terminals contact a NeuN labeled postsynaptic neuron. Primary afferents were labeled in this region of tissue but did not converge directly onto this neuron. ***D***, shows closely apposed S1 CST and primary afferent axons and terminals. CT-B is colabeled with VGLUT1 in some of the afferent terminals (light blue), which suggests they are functional. ***E***, is an additional example of the close proximity of S1 CST axons and afferent terminals (yellow circles). Inset shows a higher magnification exemplar of an individual S1 CST and primary afferent terminal in remarkably close contact. We were unable to label an interneuronal intermediate, which would be required for presynaptic inhibition (Aii), so the role of this putative connection is unclear. ***F***, shows a NeuN labeled neuron surrounded by afferent and efferent terminals. 3D assessment however showed that only the primary afferents encircled and made direct contact on to this target neuronal soma (scenario Ai or Aii).

Figures 8 and **9** show examples of some of the different ways (Figure 8A) that afferent and efferent inputs may influence each other and connect within the dorsal horn. Whilst we were unable to label dendrites or differentiate postsynaptic neuronal populations with available antibodies, we observed certain patterns which reflect circuitry described in genetically defined mouse models. We saw evidence of neurons which received strong independent input from either afferent or efferent populations (Figures 8C & 8F; may fit mechanism **8Ai**). We also saw evidence of S1 CST axons and boutons juxtaposed with afferent terminals (Figure 8B-E), which provides much of the circuitry for presynaptic inhibition (Figure 8Aii). Finally, there were convergent afferent and S1 CST inputs onto postsynaptic neuronal cell bodies (Figure 9; mechanism shown in **8Aiii**). Though we were not able to quantify any of these mechanisms, all were clearly present in both control and lesioned animals (Figures 8 & 9), and provide either a direct or indirect route for integration of sensory information within the spinal cord.

**Figure 9.**
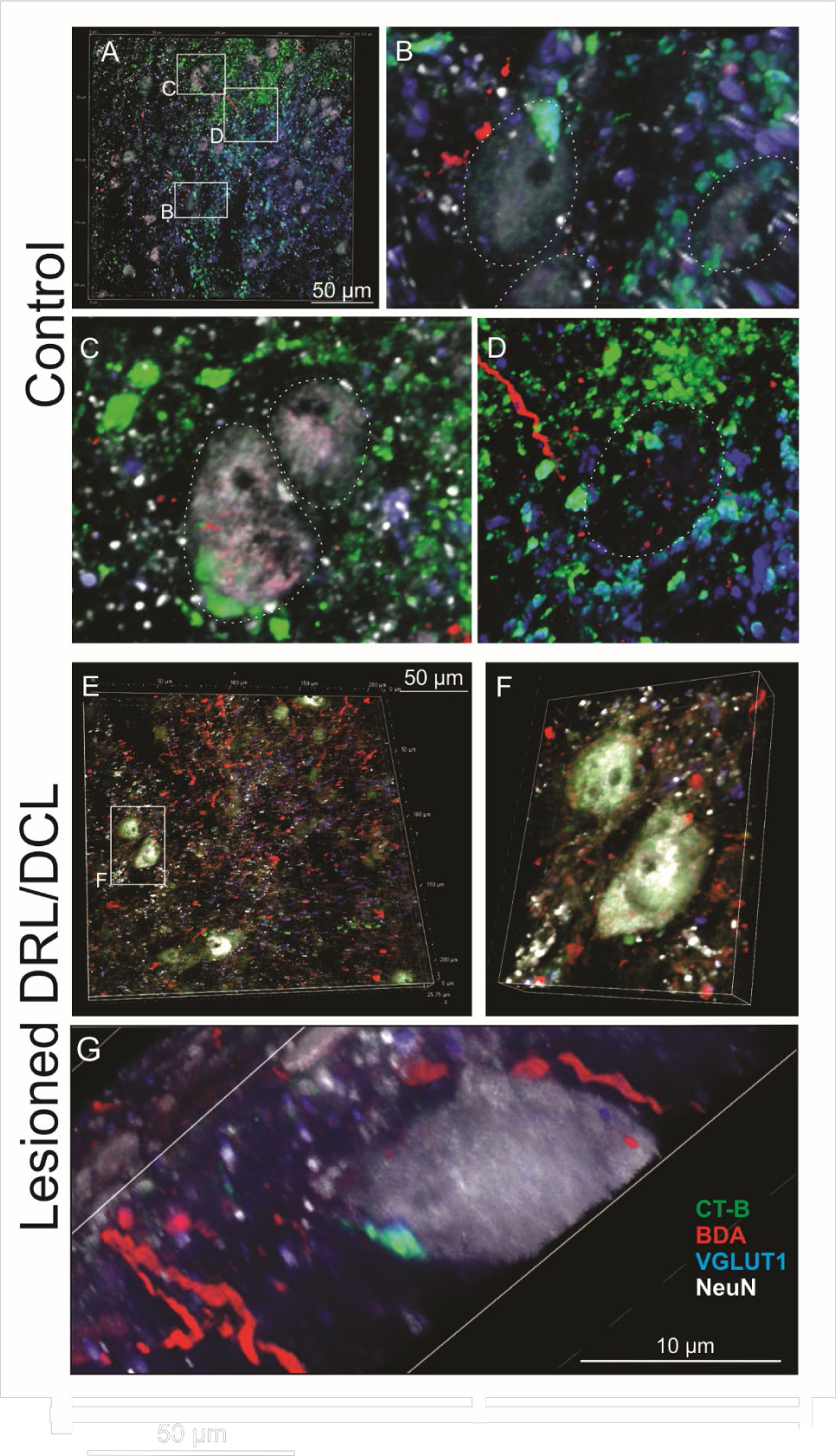
Confocal images of primary afferent (CT-B) and efferent (BDA = S1 CST) inputs onto spinal interneurons in a control (***A-D***) and lesioned monkey (***E-G***). ***A***, shows a low power image of the ventral medial dorsal horn in the control monkey 1601. Widespread colabeling (turquoise) of primary afferents (CT-B) with VGLUT1 can be seen, as well as S1 CST axons (red). ***B***, shows a NeuN+ neuron receiving convergent afferent and efferent inputs onto its cell body, which supports mechanism **Aiii** in Figure 8. ***C***, shows a neuron soma with direct input from primary afferents. ***D***, shows a NeuN-neuron covered with colabeled afferent/VGLUT1 contacts. These ‘ghost’ neurons were common and indicate that NeuN does not label all neurons. It should be noted that in the monkey, neuron and interneuron specific antibodies (e.g. NeuN, Calbindin and Parvalbumin) label only a small proportion of the total neurons present so there is a considerable amount of information missing. ***E***, shows the ventral medial dorsal horn in a lesioned monkey (1603). A greater number of CST axons (red) were evident in this region of reorganization. ***F***, shows a large NeuN+ cell receiving convergent (putative) inputs from S1 CST (red), and a primary afferent (green). ***G***, shows an example of a NeuN+ neuron with convergent S1 CST and primary afferent terminals on its cell body (also supporting **Aiii** in Figure 8).

### Compensatory effects extend along the full rostrocaudal extent of the spinal input zone

The primary afferent terminal area was diminished across the full rostrocaudal input zone following a DRL/DCL affecting digits 1-3 (Figure 10A; **top panel**). In parallel, the corresponding S1 CST terminal area increased over the same range, and particularly so rostral to the lesion (Figure 10A; **bottom panel**). This snapshot of the afferent and efferent inputs to the cord demonstrates that digit sensory information is being funnelled through many different segments in healthy animals, not just the those which are normally considered (C5-8). It also underscores a redundancy in both pathways which likely subserves co-ordination of multi-joint/postural movements in the healthy state, but which can be harnessed to promote recovery following injury.

**Figure 10.**
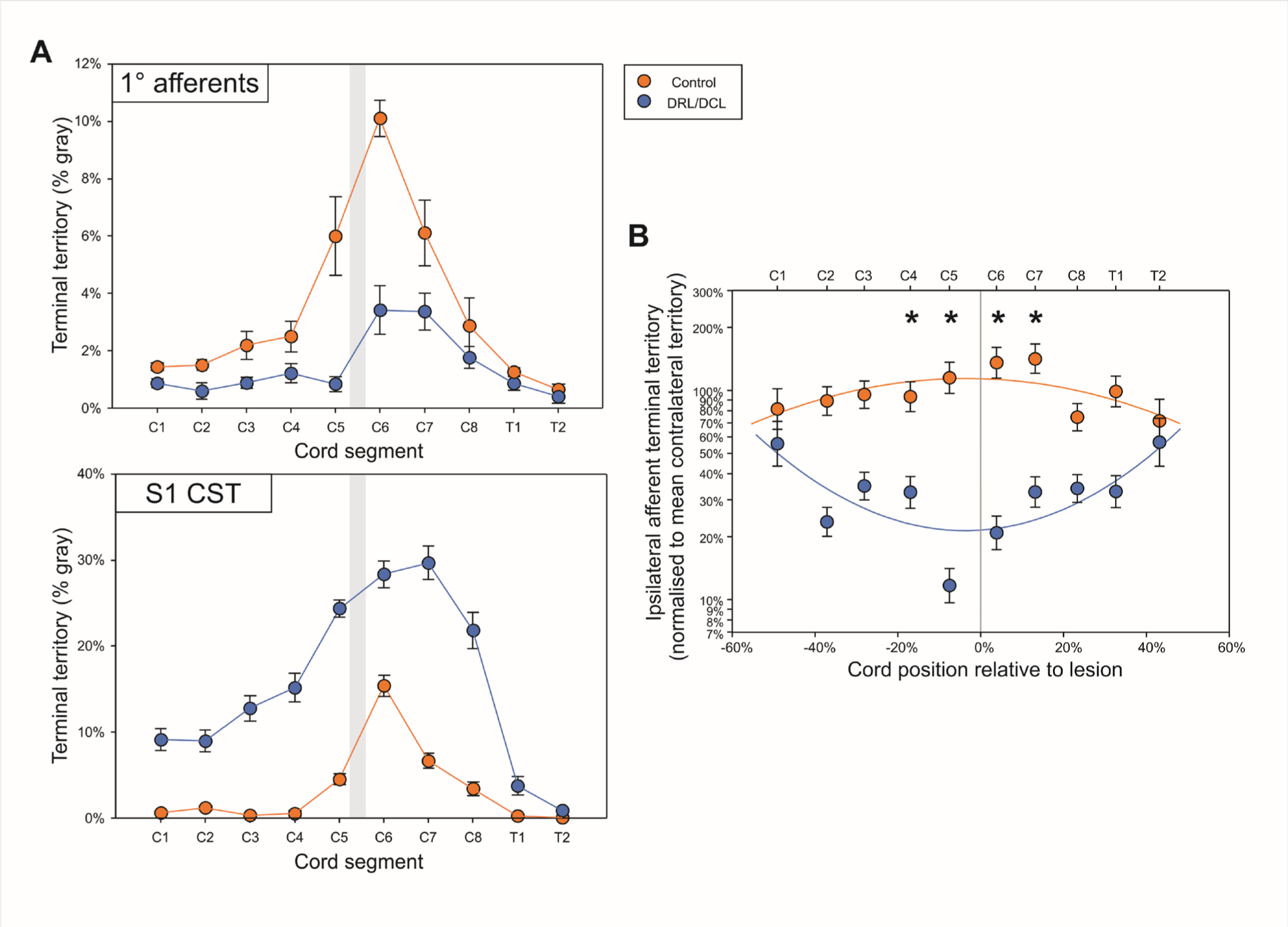
**A**, Mean rostrocaudal profiles of primary afferent (top) and S1 CST (bottom) pathways ipsilateral to the lesion (n=2 control, n=2 DRL/DCL). Error bars = SEM. Grey shaded areas indicate the lesion zone. **B**, Shows the same data as in A (top graph), but statistically analysed and presented in a different way. Here the ipsilateral afferent terminal territory was plotted along the length of the cord, as a percentage of the mean contralateral terminal territory. The position of each section was expressed relative to the lesion (to normalize position for fair comparison), with 0% indicating the location of the DCL. Data were analyzed as a log-log repeated measures polynomial regression using a REML mixed model. The corresponding contralateral terminal territory was included so that each segment acted as its own control. The ipsilateral afferent terminal territory was analyzed as a percentage of the contralateral afferent terminal territory. This showed a curvilinear relationship that differed between control and lesioned groups (F_1,113_=9.666; P=0.0024), and the corresponding least-squares line is plotted. Bonferroni-corrected post-hoc tests showed that the DRL/DCL curve (F_1,112_=7.742; P=0.0006), but not the control curve (F_1,114_=2.289; P=0.1331) differed significantly from a flat line. For ease of visualization data are plotted as the arithmetic mean (+/− SEM). Differences between the corresponding least squares means were tested with bonferroni-corrected post hoc contrasts, with significance indicated with asterisks. Control means did not differ significantly from the mean contralateral terminal territory (i.e. 100% on the graph), while DRl/DCL means differed significantly (i.e. P<0.0025) for C4, C5, C6, C7, and T1. It was not possible to analyse the S1 CST data in the same way, as contralateral territory information was not available.

### Perineuronal nets were not implicated in axonal sprouting in the dorsal horn

We used WFA to visualize perineuronal nets (PNNs) within the dorsal horn. PNNs have been implicated in regulating plasticity within CNS tissue (Sorg et al., 2016; Fawcett et al., 2019), and their digestion using the chondroitinase ABC enzyme (within the intermediate zone and ventral horn) has been linked to improved functional recovery following spinal injury (Rosenzweig et al., 2019). As illustrated in our monkeys (Figure 11), PNNs were not present in the adult dorsal horn in either normal (Figure 11A-D) or DRL/DCL animals (Figure 11E-H), though they remain abundant throughout the intermediate zone and ventral horn. This striking distribution fits with other recent work in primates (Mueller et al., 2016; Rosenszweig et al., 2019) and implies that PNNs were not important for the axonal sprouting observed in the present study. It may further suggest that the dorsal horn is a locus of ongoing plasticity/dynamic connections throughout life.

**Figure 11.**
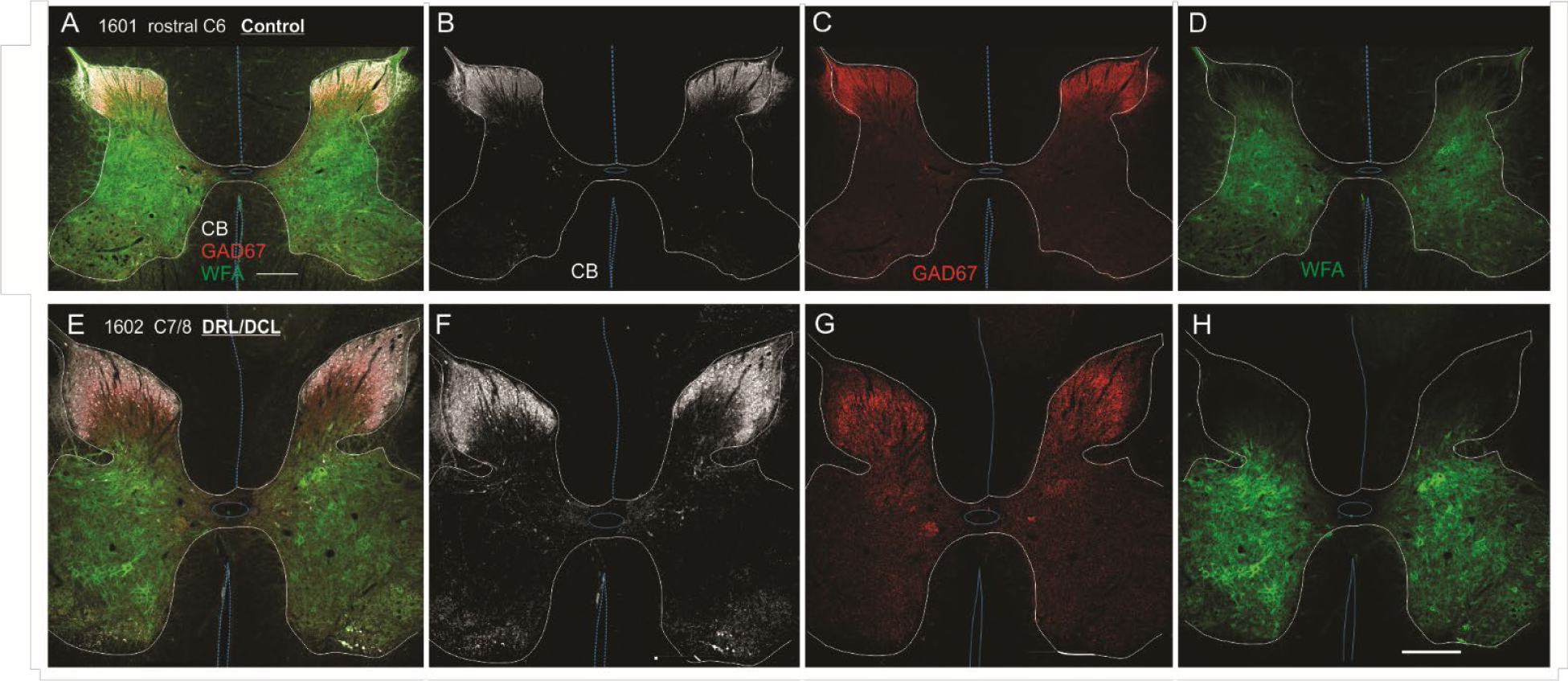
Immunofluorescence confocal images showing the regional distribution of GAD67, Calbindin, and WFA (for perineuronal nets) bilaterally in a control (**A-D**; monkey 1601) and lesioned animal (**E-H**; monkey 1602). Both sections are from the lesion zone of the DRL. The pattern of labeling shown was equivalent bilaterally, rostrocaudally, and between control and lesioned monkeys. Note that there was a distinct lack of WFA staining in the dorsal horn on either side of the cord and no degradation in either the intermediate zone or ventral horn on the side of the lesion. This implies that chondroitin sulphate proteoglycans (CSPGs) and perineuronal nets played little if any role in the S1 CST and primary afferent terminal sprouting observed in this study. Scale bar = 500µm.

### Primary afferent and S1 CST efferent bouton size distributions alter following SCI

To determine how terminal populations might alter post-lesion, we looked at the relative sizes of afferent and efferent boutons in control versus lesioned monkeys. In this analysis, a total of 660 individual boutons were measured from 26 sections taken from 4 monkeys (control =2, lesioned =2; Figure 12). Since bouton size correlates with axon caliber (Innocenti and Caminiti, 2017), the large size ranges observed in our study reflects the heterogeneity of the peripheral afferent and S1 CST populations, which both include myelinated and unmyelinated fibers of all sizes. Both afferents and efferents were found to have similar bouton size distributions in the dorsal horn (area = 0.1-9µm^2^; Figure 12A). This contrasts with the rat (Jiang et al., 2016), though experimental differences may also account for this discrepancy.

**Figure 12.**
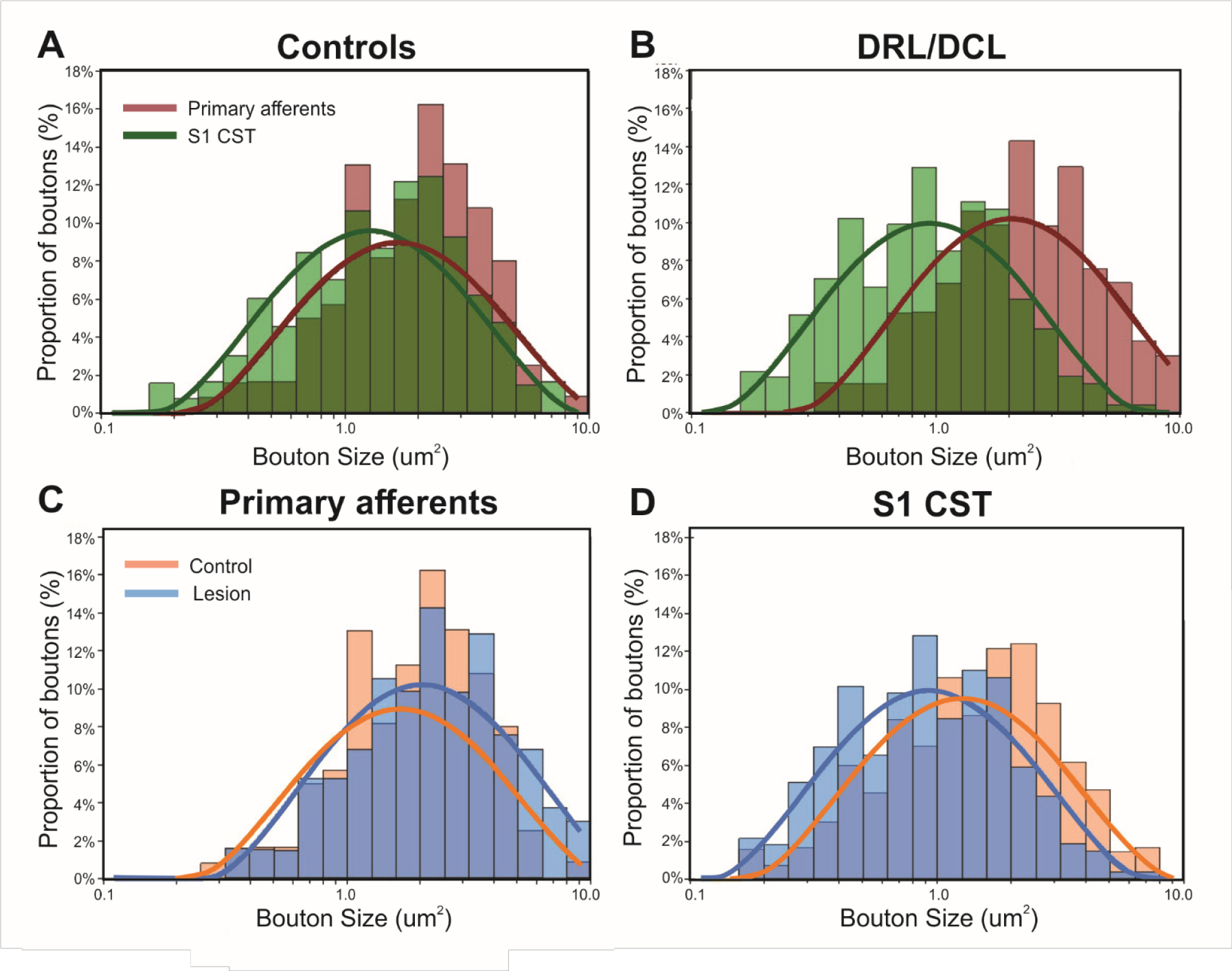
Distributional analysis of primary afferent and S1 CST bouton sizes. Data were log normal distributed, so the x-axis is plotted on a log scale. Actual (histogram) and fitted (curve) distributions are shown. Size of primary afferent and S1 CST boutons are directly compared in **A** (control animals) and **B** (DRL/DCL). This same data is then presented differently underneath to illustrate the effect of DRL/DCL on primary afferent (**C**) and S1 CST bouton size individually (**D**).

Following a DRL/DCL, there was no significant change in the primary afferent bouton size distributions, which means that boutons were lost equally across the board (Figure 12C). Since our afferent bouton profiles were equivalent between control and lesioned monkeys, CT-B uptake was not prevented in smaller fibers after injury, as has been suggested (Tong et al., 1999). For S1 CST terminals, there was a shift towards smaller boutons after DRL/DCL (Figure 12D; P = 0.0014). This indicates that even though this tract was not directly injured, it was still significantly impacted, in that there was selective loss of the largest and by inference fastest S1 CST fibers (Innocenti and Caminiti, 2017).

Our statistical analysis showed that bouton size (area) followed a lognormal distribution (Akaike information criterion, AICc = 1992) as opposed to a normal distribution (AICc = 2383) (P < 0.0001), where the model with the smallest value is the preferred model (Akaike, 1974). The shape (width) of the underlying lognormal distributions did not differ between treatment (control or lesion), bouton type (S1 CST or primary afferent) or any interaction between them. However, the peak bouton size differed for each distribution (treatment-by-bouton-type interaction: F_1,133_ = 15.88; P= 0.0001). Thus, the peak for afferent boutons was significantly lower than the peak size for S1 CST boutons for both control and lesioned animals, but this difference was significantly greater in lesioned animals (Figure 12A,B). This means that S1 CST boutons showed a significantly smaller peak size in lesioned versus control animals (Figure 12D), whereas afferent boutons did not differ significantly (Figure 12C). As such, afferent terminals were evenly eliminated by the DRL/DCL lesion, but the effect on S1 CST was more specific, causing selective loss of the largest terminals.

## Discussion

We report a number of new findings. ***First***, we show that when a DRL/DCL in monkeys removes all detectable input from the thumb, index and middle fingers (D1-D3), the small spared population of primary afferents sprout within the dorsal horn ipsilateral to the lesion. This means that in contrast to the bilateral CST response (Darian-Smith et al., 2014; Fisher et al., 2018), the central DCL does not induce afferent sprouting contralateral to the lesion. ***Second***, we show that primary afferents and S1 CST efferents terminate in an overlapping region of the dorsal horn, that extends rostrocaudally from C1 through T2. The terminal spread is far more extensive than previously reported and both populations sprout significantly through this rostrocaudal extent, begging questions about the function of these spatially distributed connections. ***Third***, we observed a number of different input strategies for primary afferents and S1 CST terminal efferents in the dorsal horn. These included independent, as well as convergent inputs on to post-synaptic target neurons and suggest a strong cortical modulation of primary afferents in normal and lesioned animals. ***Finally***, a terminal bouton size analysis showed that larger inputs from the S1 CST are lost after deafferentation, but that bouton size profiles for primary afferents do not change. We provide a functional interpretation of our findings with potential implications for recovery from injury.

### Primary afferent inputs to spinal cord

In macaques, the majority of primary afferents from the digits terminate in superficial layers of the dorsal horn within segments C5-7 (Florence et al., 1988; Brown et al., 1989; Florence et al., 1989; Florence et al., 1993). However, there is also a considerable input that terminates in the medial dorsal horn in laminae IV-VI (Figure 5), and in this investigation, this deeper input extended from C1-T2. This means that primary afferents from the digits project to a considerable portion of the spinal cord, which may provide sensory feedback more broadly for co-ordinated forelimb and body movements. The function of ‘outlying’ inputs to spinal levels rostral to those normally associated with hand function (i.e. C1-C3), is not known but they may provide some redundancy that benefits recovery after injury. The relationship between these inputs and the C3-4 propriospinal network (Isa, 2017) is also not known, though it seems reasonable to speculate that there is a connection.

Given that afferent inputs are largely absent immediately after a DRL alone (Darian-Smith, 2004), the extent of primary afferent labeling observed on the side of the lesion indicates sprouting from the tiny spared afferent population. And, even though the overall input territory to the dorsal horn remained significantly reduced at this time-point, this sprouting was considerable. Importantly, recovery has occurred at this point, so the greatly reduced afferent population, fed by a reduced number of peripheral receptors (Crowley et al., 2019), is able to support a high level of digit function (Darian-Smith and Ciferri, 2005).

### S1 CST fibers never cross the midline

Earlier work in our lab demonstrated that M1 and S1 CSTs sprout extensively on both sides of the cord following a DRL/DCL (Darian-Smith et al., 2014). Though the M1 CST is normally bilateral (Ralston and Ralston, 1985; Dum and Strick, 1996; Galea and Darian-Smith, 1997; Rosenzweig et al., 2009; Morecraft et al., 2013), S1 CST fibers are strictly contralateral in normal monkeys. Here, we show that the S1 CST remains contralateral after a DRL/DCL. The lack of midline crossing of S1 CST versus M1 CST axons after injury is likely a simple function of purpose. Movements are often bimanual or multi-jointed so fibers which span the midline are useful in driving co-ordinated output. However, a pathway which has regulatory control in triaging sensory input may need more refined access to discrete populations. There are a number of molecular factors which support the midline as a barrier to S1 CST fibers, even after injury (Hollis, 2016).

The phenotype of S1 CST fibers is quite distinct from the typically large, myelinated, fast conducting M1 CST. S1 CST axons are comparatively smaller (Ralston & Ralston, 1985) and thinly myelinated. This corresponds to smaller cell bodies in layers V and VI in S1 (Murray and Coulter, 1981), and slower conduction velocities (McComas and Wilson, 1968; Casale and Light, 1991). In general, electrical stimulation of the S1 CST does not produce movement, but does have a strong effect downstream. For example, S1 stimulation elicits large dorsal root potentials (Carpenter et al., 1963) and exerts a potent inhibitory influence in the spinal cord (Yezierski et al., 1983) and musculature (Widener and Cheney, 1997). In contrast, M1 stimulation has predominantly excitatory effects, which again highlights the stark difference between these two pathways and a differing underlying functional role for each projection.

### Integration of sensory inputs

The dorsal horn is constantly bombarded with afferent input, yet it is clear that this information is not routed to the cortex unfiltered since we do not attend to irrelevant sensory inputs. Instead, sensory inputs are integrated to streamline salient information. This occurs throughout the CNS and recently there has been much focus on the dorsal horn in the spinal cord. This region is crucial for processing and relaying sensory information, yet very little is known about the detailed circuitry, particularly in primates. Recent genetic studies in mice report at least 11 different interneuron types in just laminae III-IV (Del Barrio et al., 2013; Bourane et al., 2015; Abraira et al., 2017), and many of these types are located in the region of convergence for sensory inputs from the CST and peripheral fibers.

In this study, the terminal territory of the S1 CST overlapped almost completely with primary afferent inputs, and this was true from C1-T2. This spatial proximity supports a role in the integration and/or gating of sensory information. There are a number of potential mechanisms that could account for, or contribute to, sensory integration. It could be achieved by specialized neurons which receive independently relayed inputs from dorsal horn interneurons (Figure 8 Ai, C, F). Convergence may also represent a mechanism by which sensory integration could occur, with the potential for S1 CST fibers to exert direct influence over sensory inputs, as shown recently in the rodent spinal cord (Hantman and Jessell, 2010; Liu et al., 2018). We observed examples of converging primary afferent and S1 CST efferent terminals on the same neurons in the dorsal horn (Figure 9). We could not identify the target cells as interneurons, but they share similarities with RORα interneurons in mice which receive mechanoreceptor as well as CST inputs (Bui et al., 2015).

#### Cortical influence over sensory inputs

The modulatory effect of the CST on afferent input has been examined for decades, though most of this work pertains to the M1 CST (Carpenter et al., 1962; Rudomin, 1999). It is known that presynaptic inhibition can selectively suppress afferent inputs to spinal interneurons during movement, presumably to retain the fidelity of motor commands (Seki et al., 2003). This has been demonstrated via increased sensory thresholds (Chapman et al., 1987; Williams and Chapman, 2000), reduced activity in the dorsal column pathway (Ghez and Lenzi, 1971), and suppression of somatosensory evoked potentials (Rushton et al., 1981; Chapman et al., 1988; Morita et al., 1998; Seki and Fetz, 2012) during movement execution. Our observation of closely apposed S1 CST and primary afferent axons and boutons in the dorsal horn (Figure 8D-E) may indicate there are similar mechanisms at play for top down control of sensory inputs by somatosensory cortex, but we cannot conclusively demonstrate this with our current dataset.

### How is sensory integration affected by injury?

We do not yet fully understand how sensory integration is impacted by injury. DRL/DCL animals appear to have normally functioning digits 2-3 months post injury (Figure 3, 4), yet it is difficult to determine whether sensory perception has returned to normal or whether they experience hypersensitivity of the affected fingers. We know that Meissner’s Corpuscles, which reduce in number over the months following a DRL/DCL, repopulate within a year post-lesion (Crowley et al., 2019), but we don’t know whether their functionality could be considered normal. In addition, little is known about the functionality of the afferent or efferent sprouts at any of the C1-T2 segmental levels, or whether this growth is useful or maladaptive (Florence et al., 1994; Koerber et al., 1994).

Sensory gating is an adaptive process which is capable of compensating for alterations in input signal. However, it is unlikely that there is a simple corrective gain change following injury given the complexity of the sensorimotor apparatus impacted by a lesion and the lengthy recovery time. The impact on integration of sensory information in general must be considerable, particularly given the compensatory strategies monkeys use to perform the retrieval task over the long term.

Local changes in inhibitory connections are likely to be important post-lesion. Decreased inhibitory transmission, and presynaptic inhibition in the dorsal horn have been reported (Calancie et al., 1993; Castro-Lopes et al., 1993; Zhang et al., 1994; Moore et al., 2002; Meisner et al., 2010), though the process is poorly understood and clearly dynamic (Darian-Smith et al., 2010).

More widespread changes also have relevance since the effects of a DRL/DCL are not restricted to local neuronal populations. The brainstem, thalamus, and cerebellum, as well as the somatosensory and motor cortex (Evarts and Tanji, 1974; Lemon and Porter, 1976), all receive afferent input, and compensatory adaptations following SCI are common (Darian-Smith and Fisher, 2019). Keeping sight of the distributed nature of recovery is important, both for understanding the reaches of post-injury reorganization, and for developing therapeutic strategies. In addition, the post-injury time point at which experimental observations are made is only a snapshot of a sequence of changes. To this end, we have yet to determine if (and to what extent) the sprouting observed 4-5 months post injury in this study is pruned (or not) in the chronic state. Transient compensatory strategies have been demonstrated after stroke (Marshall et al., 2000; Ward et al., 2003, 2004; Dancause et al., 2015) and in SCI (Nishimura et al., 2007; Isa, 2017), so determining widespread circuit remodelling during the chronic phase will be especially important and help predict recovery outcome.

The findings of the present study add important new insights into how this region of sensory convergence adapts following even a small spinal deafferentation injury. However, our findings barely scratch the surface and we are a long way from confidently identifying the key changes that directly lead to functional recovery. Understanding the changing dorsal horn circuitry, within the broader sensorimotor system is clearly important, and is therefore critical for the informed development of targeted rehabilitative therapies for spinal injuries.

## Acknowledgements

This work was supported by the National Institute of Neurological Disorders and Stroke (R01 NS091031 to CD-S). Thanks also to Alayna Lilak for her technical help with the monkeys, and Matthew Crowley.

